# Missense Mutation ADNP p.C687R Disrupts Chromatin Regulation and GABAergic Differentiation in HVDAS

**DOI:** 10.1101/2025.11.10.687585

**Authors:** Qi Chen, Xixi Liu, Mengnan Wu, Daijing Sun, Xingyu Ding, Mengling Zhou, Wenzhu Peng, Yan Cheng, Biqing Xue, Ning Tang, Gang Xu, Yilin Tai, Qiong Xu, Man Xiong, Yan Jiang

**Affiliations:** Institutes of Brain Science, State Key Laboratory of Medical Neurobiology and MOE Frontiers Center for Brain Science, Fudan University, Shanghai 200032, China; Department of Child Health Care, Children’s Hospital of Fudan University, Shanghai 201102, China; Multiscale Research Institute of Complex Systems, Fudan University, Shanghai 200433, China

**Keywords:** ADNP, GABAergic neural progenitors, gain-of-function, Helsmoortel–Van der Aa syndrome, histone modification

## Abstract

Activity-dependent neuroprotective protein (ADNP) is a key regulator of neurodevelopment and a high-risk gene for autism spectrum disorder (ASD). Most pathogenic variants cause loss-of-function in Helsmoortel–Van der Aa syndrome (HVDAS), yet the impact of missense mutations remains unclear. Here, we characterized a ADNP missense mutation (p.C687R) identified from a HVDAS patient, predicted to disrupt its ninth zinc finger domain. Overexpression of p.C687R altered subnuclear localization and affected wild-type ADNP distribution. *In utero* electroporation revealed impaired neuronal migration and cortical branching. CUT&Tag profiling showed mutant-specific chromatin binding, preferentially targeting histone modification genes through both direct and distal interactions. Patient-derived iPSCs carrying p.C687R exhibited upregulated bivalent histone marks (H3K4me3 and H3K27me3) on neurodevelopmental genes, including regulators of GABAergic differentiation, which became transcriptionally activated in neural progenitors. These findings reveal a gain-of-function mechanism by which ADNP p.C687R perturbs chromatin regulation and disrupts GABAergic lineage specification, expanding the molecular framework of ADNP pathogenesis.

**The Paper Explained Problem:** Mutations in the gene ADNP cause Helsmoortel–Van der Aa syndrome (HVDAS), a severe neurodevelopmental disorder, and also increase the risk for autism spectrum disorder (ASD). While most known ADNP mutations lead to loss of protein function, many missense mutations that alter a single amino acid remain poorly understood regarding their effects on brain development and disease contribution.

**Results:** We characterized a specific missense mutation, ADNP p.C687R, from a patient diagnosed with HVDAS, and found that it does not simply inactivate the protein but instead confers a disruptive new function. Across multiple models, from cells to mice, the p.C687R mutant exhibits abnormal chromatin binding, altered subnuclear localization, and impaired neuronal migration and differentiation. Importantly, in patient-derived iPSCs, it marks distinct neurodevelopmental genes with bivalent histone modifications, which become aberrantly activated during differentiation into GABAergic neural progenitor cells.

**Impact:** These findings confirm the pathogenicity of the missense mutation ADNP p.C687R in HVDAS, reveal a gain-of-function mechanism whereby the mutant disrupts chromatin regulation and GABAergic lineage specification, and advance our understanding of ADNP-related neurodevelopmental disorders.

## Introduction

Helsmoortel-Van der Aa syndrome (HVDAS; OMIM 615873), also known as ADNP syndrome, is a rare autosomal dominant neurodevelopmental disorder characterized by core features such as developmental delay, intellectual disability, autism spectrum disorder (ASD), hypotonia, and sensory processing abnormalities. Additional comorbidities may involve dermatologic, respiratory, gastrointestinal, and urogenital systems (Pascolini *et al*, 2024). HVDAS is caused by pathogenic variants in the *ADNP* gene and is estimated to account for approximately 0.17% of all ASD cases (Helsmoortel *et al*, 2014).

*ADNP* encodes activity-dependent neuroprotective protein (ADNP), a highly conserved transcriptional regulator essential for neurodevelopmental processes, including neural tube closure (Pinhasov *et al*, 2003), cortical patterning (Clemot-Dupont *et al*, 2025), axonal outgrowth (Chen & Charness, 2008), and synaptic plasticity (Cho *et al*, 2023). ADNP was first identified via its N-terminal neuroprotective peptide NAPVSIPQ (NAP) (Bassan *et al*, 1999), which promotes neuronal survival (Zemlyak *et al*, 2000). Structurally, ADNP contains nine zinc finger (ZF) domains, a nuclear localization signal (NLS), and a C- terminal homeobox domain, enabling its function in gene regulation and chromatin remodeling (Yan *et al*, 2022). It is a critical component of the ChAHP complex (CHD4/ADNP/HP1), which modulates chromatin accessibility and 3D genome architecture (Kaaij *et al*, 2019).

Since the first identification of *ADNP* variants in HVDAS patients in 2014 (Helsmoortel *et al*., 2014), several cohorts have defined the genetic and phenotypic spectrum of the disorder (Bend *et al*, 2019; Ge *et al*, 2024; Pascolini *et al*., 2024; Siper *et al*, 2021; Van Dijck *et al*, 2019). The majority of pathogenic variants are *de novo* coding mutations, most commonly nonsense and frameshift changes that introduce premature termination codons (PTCs), leading to truncated proteins. A large-scale study, including a cohort of 78 patients from 16 countries (Van Dijck *et al*., 2019), and additional case series across Europe (Pascolini *et al*., 2024), North America (Bend *et al*., 2019; Siper *et al*., 2021), and Asia (Ge *et al*., 2024), have consistently shown that truncating variants dominate the ADNP mutational landscape. Notably, recurrent variants such as p.Tyr719*, p.Arg730*, and p.Leu832Ilefs*81 have been observed in multiple unrelated individuals (Bend *et al*., 2019; Ge *et al*., 2024; Siper *et al*., 2021; Van Dijck *et al*., 2019).

Mechanistic studies to date have also primarily focused on truncating variants (Cappuyns *et al*, 2018). The loss of key functional domains, particularly the NLS and homeobox, leads to impaired nuclear import and disrupted chromatin binding (Cappuyns *et al*., 2018; Ganaiem *et al*, 2022; Yan *et al*., 2022). For instance, variants affecting residues 473–719 result in cytoplasmic retention (Cappuyns *et al*., 2018), and p.Tyr719* is associated with more severe motor and sensory deficits (Van Dijck *et al*., 2019). Recent work has also linked variant position to distinct genome-wide DNA methylation signatures (Bend *et al*., 2019) and clinical severity (Van Dijck *et al*., 2019), suggesting genotype-phenotype correlations. Functional studies of truncating ADNP variants have showed impaired functions in R-looping (Yan *et al*., 2022), Wnt/β-catenin signaling (Sun *et al*, 2020a), microtubule dynamics (Ivashko-Pachima *et al*, 2017), and autophagy regulation (Merenlender-Wagner *et al*, 2015). These deficits are recapitulated in mouse models including haploinsufficient (Cho *et al*., 2023), brain conditional knockout (Clemot-Dupont *et al*., 2025), and p.Tyr718* knock-in mice (Shapira *et al*, 2025), supporting a loss-of-function (LOF) mechanism in HVDAS pathogenesis.

In contrast, the clinical relevance and functional impact of ADNP missense variants remain poorly understood. While several missense variants have been reported in databases, few have been linked to HVDAS or studied in detail. Among five missense variants analyzed in a 2019 study, only one (p.Gln67His) was *de novo* and classified as likely pathogenic (Bend *et al*., 2019). The first validated disease-associated missense variant disrupted the terminal residue of the NAP peptide in a patient with mild neurodevelopmental symptoms (Gozes & Shazman, 2023). Our previous work identified two additional *de novo* missense variants, p.C687R and p.R730G, in a Chinese pediatric cohort (Ge *et al*., 2024), yet their pathogenic mechanisms remain to be explored.

In this study, we focus on the p.C687R variant to investigate its functional consequences and underlying molecular mechanism. This variant lies near the ninth ZF domain and is predicted to disrupt ZF structure, potentially altering the DNA-binding capacity of ADNP. To evaluate its impact, we used overexpression cell models to assess the nuclear localization, chromatin binding, and gene regulatory functions of ADNP (p.C687R), and conducted *in utero* electroporation (IUE) in mouse embryos to examine the effects on neuronal differentiation. Moreover, we generated patient-derived iPSCs and differentiated them into GABAergic neural progenitor cells (NPCs) and performed ChIP-seq, ATAC-seq and RNA-seq to profile H3K4me3 and H3K27me3 landscapes, chromatin accessibility, and gene expression, thereby exploring how p.C687R affects neurodevelopmental programs, particularly GABAergic neuron specification. Our findings suggest that p.C687R may exert gain-of-function effects, expanding the current understanding of HVDAS pathogenesis beyond the classical LOF model associated with truncating variants.

## Results

### Characterization of *ADNP* variants with the focus on missense mutations

To systematically investigate the distribution and potential clinical relevance of *ADNP* variants, we compiled and analyzed both previously published and newly collected cases. Previously reported cohorts included 78 cases from Belgium (Van Dijck *et al*., 2019), 22 from Canada (Bend *et al*., 2019); 22 from the United States (Siper *et al*., 2021); and 15 from our earlier study (Ge *et al*., 2024), total 137 cases with confirmed *ADNP* variants and clinical diagnoses. These cases were classified as the ‘Clinical’ group. Additionally, we newly collected 74 cases identified through peripheral blood exome sequencing in China (42 males and 32 females), and retrieved 802 *ADNP* variant entries from the NCBI ClinVar database (https://www.ncbi.nlm.nih.gov/clinvar/, updated as of June 23, 2025). Since these 876 cases lacked confirmed clinical diagnoses, we classified them as the ‘Non-Clinical’ group.

We first analyzed the mutation types across both groups **(Fig. 1A)**. Among the 137 ‘Clinical’ cases, the majority of mutations were nonsense (N = 72, 52.6%) or frameshift (N = 60, 43.8%), consistent with the known LOF mechanism of ADNP pathogenesis. Only four missense variants were identified in this group, which are: ADNP (p.M1V), a start codon mutation; ADNP (p.Q67H), located before the first ZF domain on the N’-end; ADNP (p.C687R), located immediately downstream of ZF domain #9; and ADNP (p.R730G), located in the NLS domain. In contrast, in ‘Non-Clinical’ group, missense (N = 492, 56.2%) were the most common, followed by synonymous (N = 164, 18.7%), frameshift (N = 108, 12.3%), and nonsense (N = 44, 5%). We then assessed the predicated pathogenicity of all variants. As expected, 97.8% of variants in the ‘Clinical’ group were classified as pathogenic or likely pathogenic (P/LP), while only 19.2% of ‘Non-Clinical’ fell into this category. Notably, 45.2% of the ‘Non-Clinical’ variants were categorized as variants with uncertain significance (VUS) **(Fig. S1A)**. Focusing specifically on missense variants, all four ‘Clinical Missense’ variants were classified as P/LP. In contrast, among the ‘Non-Clinical Missense’ variants, 69.7% were VUS, 18.9% were benign or likely benign (B/LB), and only 1.6% were P/LP **(Fig. 1B)**. These findings reveal a major gap in interpreting ADNP missense variants, as most remain of uncertain significance in undiagnosed cases. Prior studies have found that the distribution of *ADNP* variants correlates with genome-wide DNA methylation patterns (Bend *et al*., 2019). As shown in **Fig. 1C**, variants located within the c.2000-2340 region (corresponding to p.667-780, which includes the NLS and harbors several high-frequency variants) are associated with global hypermethylation and are classified as ‘epi-ADNP-2’. In contrast, variants outside this region exhibit global hypomethylation and are classified as ‘epi-ADNP-1’ (Bend *et al*., 2019). Among the 136 ‘Clinical’ cases, 54% (N = 73) fell into the ‘epi-ADNP-1’ group and 46% (N = 63) to the ‘epi-ADNP-2’ group. In ‘Non-Clinical’, the vast majority (N = 561, 85%) fell into the ‘epi- ADNP-1’ group, while only 15% (N = 101) were located in epi-ADNP-2 **(Fig. 1C, Fig. S1B)**. To date, only one pathogenic missense variant in coding sequence (CDS) of ADNP, ADNP (p.Q362E), has been reported, which was associated with mild developmental delay (Gozes & Shazman, 2023). In our earlier work, we reported two CDS missense variants, including a *de novo* variant, ADNP (p.C687R), located just one amino acid downstream of the #9 ZF domain. The patient carrying this variant exhibited core clinical features consistent with the diagnosis of HVDAS, including global developmental delay in early childhood, subsequent intellectual disability, autistic traits, distinctive facial dysmorphism, attention deficits, sleep disturbances, hypotonia, and multisystem involvement affecting the skeletal and urinary systems. She showed short stature before 5 years of age, followed by catch-up growth. Brain MRI demonstrated mild ventricular enlargement and minimal mastoid T2 hyperintensity, while EEG findings were unremarkable **(Table 1**, see **Supplement Material** for detailed case report). In this study, we focused on hADNP (p.C687R) to investigate its pathogenicity. Protein 3D structure prediction revealed that the mutant hADNP (p.C687R) adopts a more compact conformation compared to the wild-type hADNP **(Fig. S1C)**. Notably, in the 601–700 aa region encompassing #8 and #9 ZF domains, four residues were predicted to be structurally affected by the p.C687R substitution. Among them, R688 and M675 showed the highest predicted impact, likely contributing to alterations in the α-helical structure of #9 ZF domain **(Fig. 1D)**. Based on these predictive results, we hypothesized that hADNP (p.C687R) may impair the function of ADNP as a transcription factor (TF) by disrupting its ZF conformation and altering genomic binding profile, thus contributing to disease onset.

**Figure 1.**
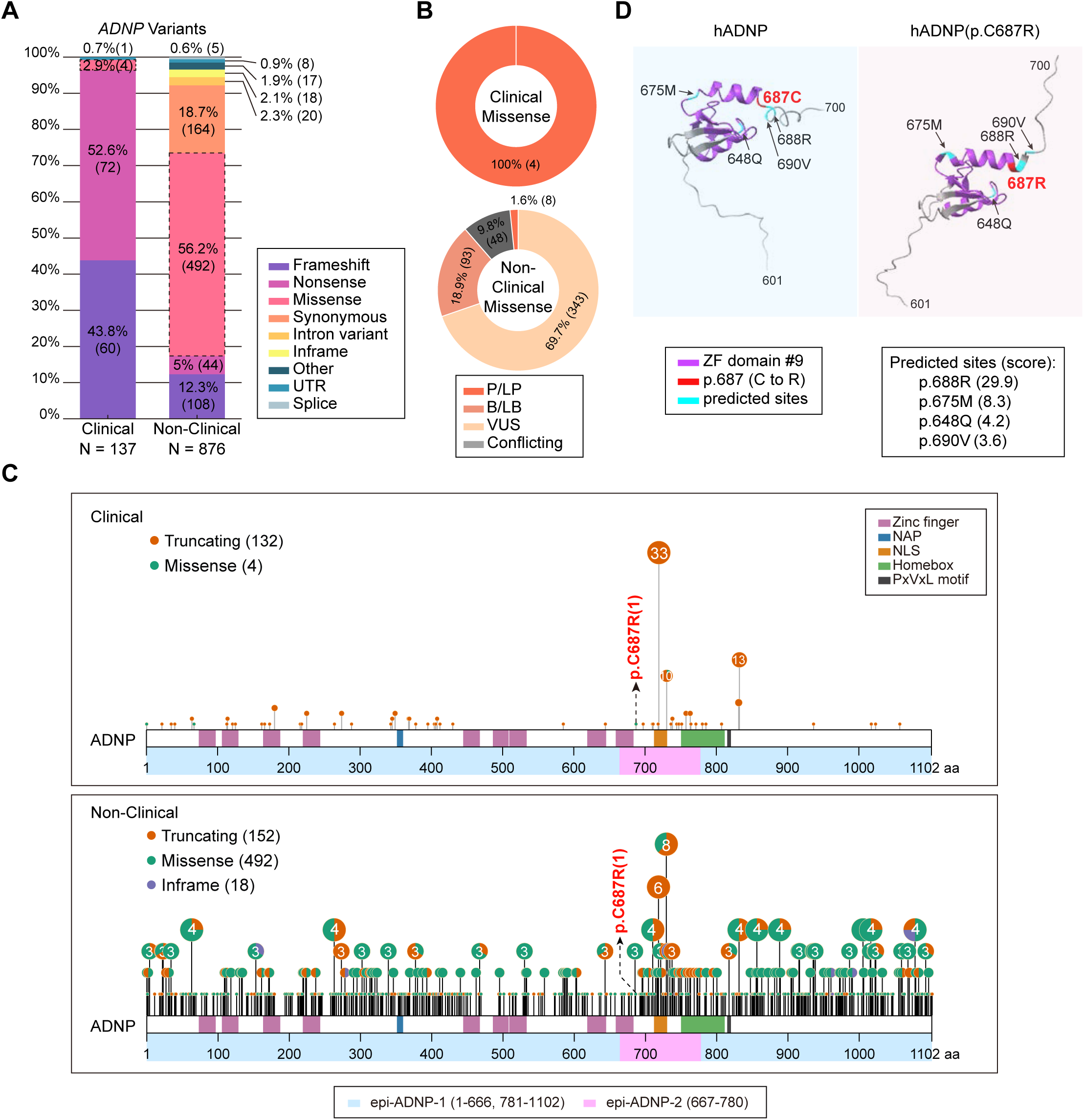
Classification of *ADNP* variants in clinical and non-clinical cohorts. (**A**) Variant classification: clinical (N = 137) vs. non-clinical (N = 876). (**B**) Predicted pathogenicity of missense variants: clinical (N = 4) vs. non-clinical (N = 492). P/LP, pathogenic/likely pathogenic; B/LB, benign/likely benign; VUS, variants of uncertain significance; Conflicting, conflicting interpretations. (**C**) Schematic distribution of variants across *ADNP* coding sequence (CDS). Upper, clinical (N = 136, one UTR variant excluded); Lower, non-clinical (N = 662, synonymous, in-frame, non-coding, and other variants excluded). Truncating variants include nonsense and frameshift variants. Blue/pink squares indicate epi-ADNP-1/2. p.C687R highlighted in red. (**D**) Predicted structural impacts of p.C687R on zinc finger (ZF) domain #9. Purple, ZF domain #9; red, mutated residue; blue, affected residues with predicted impact scores.

**Table 1:** Clinical phenotypic analysis of the patient carrying ADNP (p.C687R) variant.

| Patient information |  |
| --- | --- |
| Age | 9 years |
| Gender | Female |
| Clinical symptoms |  |
| Global developmental delay/Intellectual disability | + |
| Autistic traits | + |
| ADHD | + |
| Sleep disturbance | + |
| Distinctive facial dysmorphism | + |
| Hypotonia | + |
| Pes planus | + |
| Multisystem involvement (skeletal and urinary systems) | + |
| Short stature | - |
| Auxiliary examinations |  |
| Brain MRI abnormalities | + |
| EEG abnormalities | - |
| Echocardiographic abnormalities | - |

### hADNP (p.C687R) alters nuclear puncta formation in HEK293T cells

ADNP is predominantly localized in the nucleus, and previous studies, including our own, have shown that pathogenic mutants can alter its nuclear localization (Cappuyns *et al*., 2018; Ge *et al*., 2024), potentially contributing to disease. To investigate this, we examined the nuclear localization of hADNP (p.C687R) in HEK293T cells and assessed its effect on wild-type hADNP under co-expression, modeling the heterozygous state in patients. Cells were transfected with hADNP, hADNP (p.C687R), or both **(Fig. 2, Fig. S2)**. In all conditions, both proteins localized to euchromatic nuclear regions, consistent with prior reports (Ostapcuk *et al*, 2018). However, their intranuclear distribution differed: wild-type hADNP formed numerous small puncta, whereas hADNP (p.C687R) was largely diffusely distributed in the nucleus and lacked puncta, and when present, puncta were larger **(Fig. 2A)**. Co-expression led to strong co-localization and normalized puncta characteristics, including the proportion of cells with puncta **(Fig. 2B)**, puncta number per cell **(Fig. 2C-D)**, and size **(Fig. 2E)**. Notably, in large puncta, hADNP signals concentrated at the periphery, while hADNP (p.C687R) signals were more uniformly distributed **(Fig. 2A)**.

**Figure 2.**
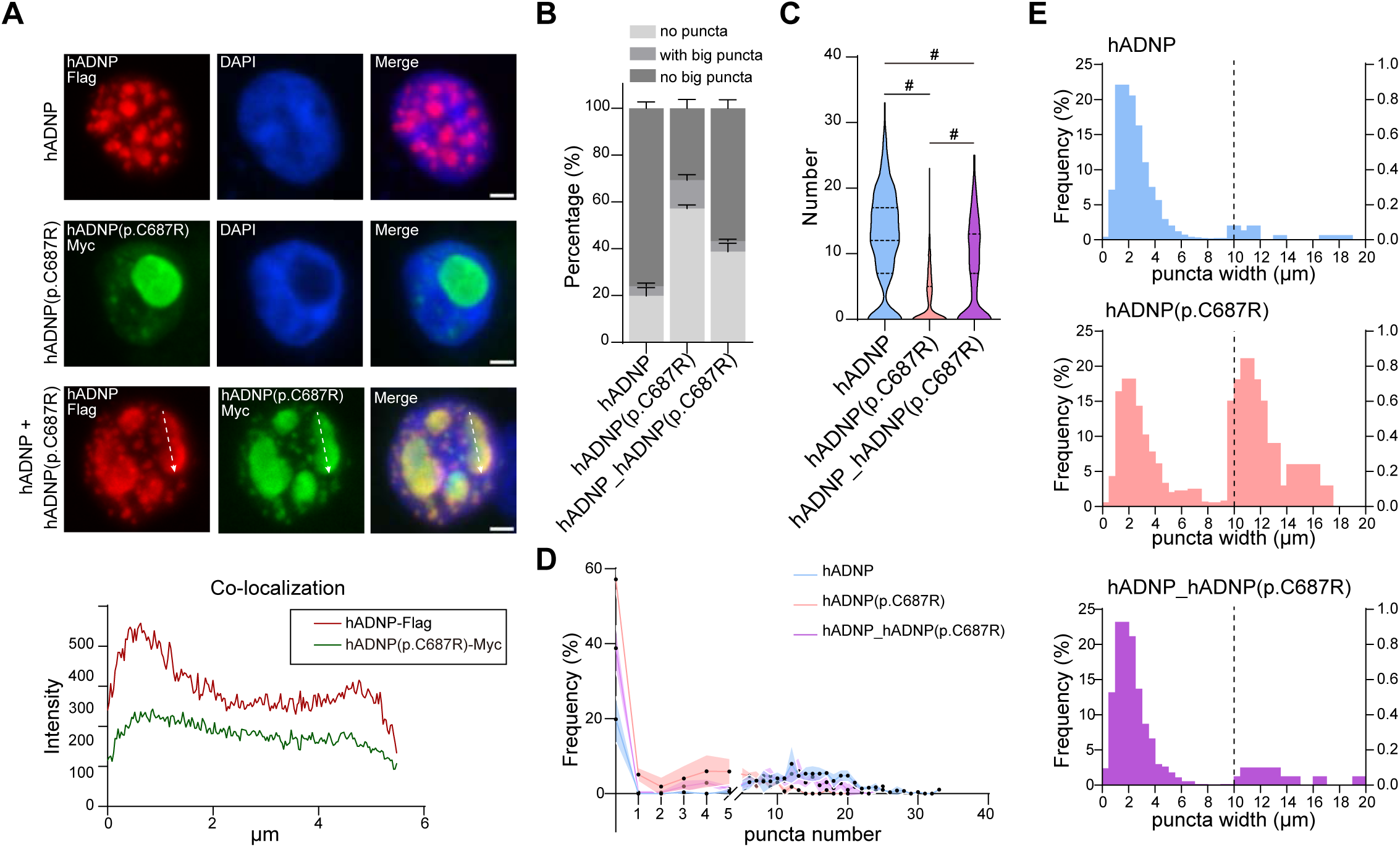
Nuclear localization of hADNP and hADNP (p.C687R) in HEK293T cells. (**A**) Top, representative images of cells transfected with hADNP-Flag, hADNP (p.C687R)-Myc, or co-transfection of both plasmids. Red, hADNP-Flag. Green, hADNP (p.C687R)-Myc. Blue, DAPI. Scale bars, 2μm. Bottom, co-localization analysis. White dashed lines indicate measured paths. (**B**) Proportion of cells with different puncta patterns: no puncta, large (>10 μm), small (<10 μm) (Mean ± SEM, N = 3/group). (**C**) Violin plots show the number of puncta per cell (Mean ± SEM, N = 3/group). Two-tailed Mann-Whitney U test, ^#^ *P* < 0.0001. (**D**) Distribution of puncta number per cell (Mean ± SEM, N = 3/group). (**E**) Puncta width (>10 μm or <10 μm) frequency.

Quantitative analysis confirmed these observations. Only 20% of hADNP-transfected cells lacked puncta, compared to nearly 60% of hADNP (p.C687R) cells; co-expression reduced this close to 40% **(Fig. 2B)**. Violin plots showed that hADNP (p.C687R) significantly reduced the average puncta number, partially rescued by co-expression **(Fig. 2C)**. Puncta number per cell was typically 5 – 25 for hADNP, 1 – 10 for hADNP (p.C687R), and intermediate for co-transfected cells **(Fig. 2D)**. Using 10 μm as a cutoff, puncta in hADNP cells were predominantly small (< 10 μm) with very few large ones; whereas hADNP (p.C687R) cells showed a marked increase in large puncta (> 10 μm); co-transfection restored the size distribution toward wild-type levels **(Fig. 2E)**.

Given the high conservation of ADNP between human and mouse, we repeated these experiments with mouse homologs mADNP and mADNP (p.C686R) in HEK293T cells **(Fig. S3A)**. While co-localization intensity **(Fig. S3B)** and puncta width differences **(Fig. S3F)** were less pronounced than in human ADNP, puncta number and distribution patterns **(Fig. S3C-E)** were consistent with human findings.

### hADNP (p.C687R) impairs neuronal migration and differentiation *in vivo*

To investigate the impact of hADNP (p.C687R) on neuronal development *in vivo*, we performed IUE at embryonic day 14.5 (E14.5), a critical stage of cortical neurogenesis when neurons are actively generated and migrate toward the cortical plate. Plasmids expressing either wild-type hADNP or hADNP (p.C687R) were co-electroporated with a GFP reporter into the lateral ventricles of CD-1 mouse embryos, and brains were examined at E18.5 by immunofluorescence staining **(Fig. 3A)**. In embryos expressing wild-type hADNP, most GFP⁺ neurons successfully migrated into the cortical plate. In contrast, a significant fraction of hADNP (p.C687R)⁺ neurons remained abnormally within the deep layer, indicating impaired radial migration **(Fig. 3A)**. Among neurons that reached the cortical plate, hADNP⁺ neurons exhibited long, slender axons typical of differentiating cortical pyramidal neurons, whereas hADNP (p.C687R)⁺ neurons displayed shortened or absent axonal projections **(Fig. 3A, inserted images)**.

**Figure 3.**
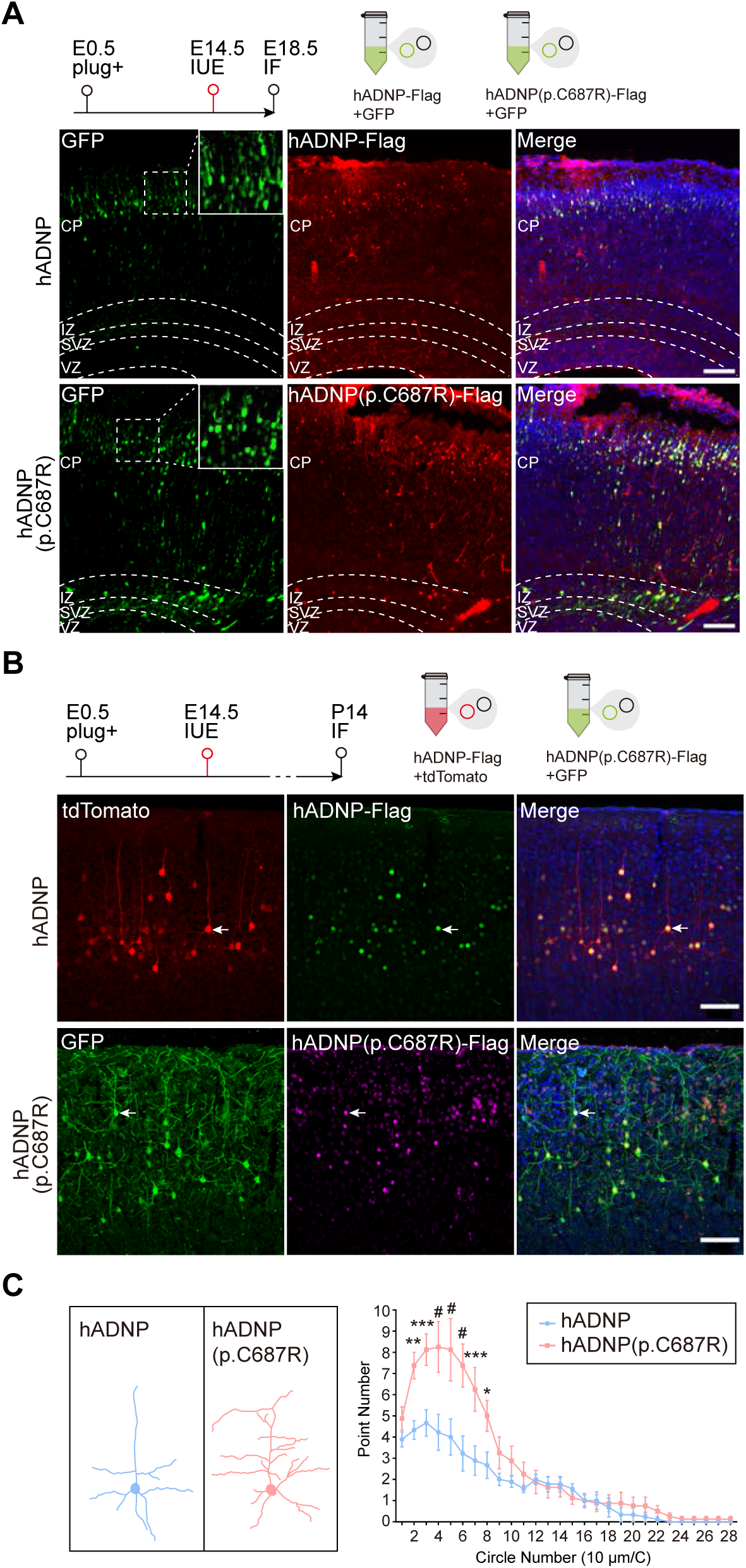
hADNP (p.C687R) impairs neuronal migration and differentiation *in vivo*. (**A**) Representative images at E18.5 after mouse cortial *in utero* electroporation (IUE) at E14.5 with hADNP-Flag + GFP or hADNP (p.C687R)-Flag + GFP. Red, Flag; green, GFP; blue, DAPI. Scale bar, 100 μm. (**B**) Representative images at P14 of IUE at E14.5 with hADNP-Flag + tdTomato or hADNP (p.C687R)-Flag + GFP showing dendritic complexity. hADNP: tdTomato (red), Flag (green); hADNP (p.C687R): GFP (green), Flag (magenta); DAPI (blue). Scale bar, 100 μm. (**C**) Sholl analysis quantifying dendritic complexity (Mean ± SEM; N = 8-9 cells/2 animals). Two-way ANOVA with post hoc, * *P* < 0.05, ** *P* < 0.01, *** *P* < 0.001, ^#^ *P* < 0.0001.

To further evaluate postnatal neuronal morphology, hADNP and hADNP (p.C687R) were co-electroporated with tdTomato and GFP, respectively, into E14.5 embryos **(Fig. 3B)**. At postnatal day 14 (P14), wild-type hADNP⁺ neurons exhibited typical mature pyramidal morphology with a single apical dendrite projecting toward the pia and well-organized basal dendrites. In contrast, hADNP (p.C687R)⁺ neurons showed markedly increased dendritic complexity, particularly near the soma **(Fig. 3B-C)**.

Together, these results indicate that the hADNP (p.C687R) mutant can disrupt neuronal migration and differentiation during mouse cortical development.

### Altered chromatin binding profile of hADNP (p.C687R) in HEK293T cells

As shown in **Fig. 1D**, protein structure prediction indicates that the p.C687R mutation may disrupt the ZF structure of hADNP, which is critical for genomic binding and affect gene regulation. To investigate whether this contributes to the observed cellular phenotypes, we overexpressed wild-type hADNP or hADNP (p.C687R), tagged with Flag at the C-terminal and HA at the N-terminal, in HEK293T cells, alongside an EGFP control. Western blotting using anti-ADNP revealed low endogenous ADNP expression in untransfected negative control (NC) or EGFP-transfected cells, whereas both hADNP and hADNP (p.C687R) were robustly expressed as full-length proteins, as validated by anti-Flag and anti-HA antibodies **(Fig. 4A, Fig. S4A-B)**.

**Figure 4.**
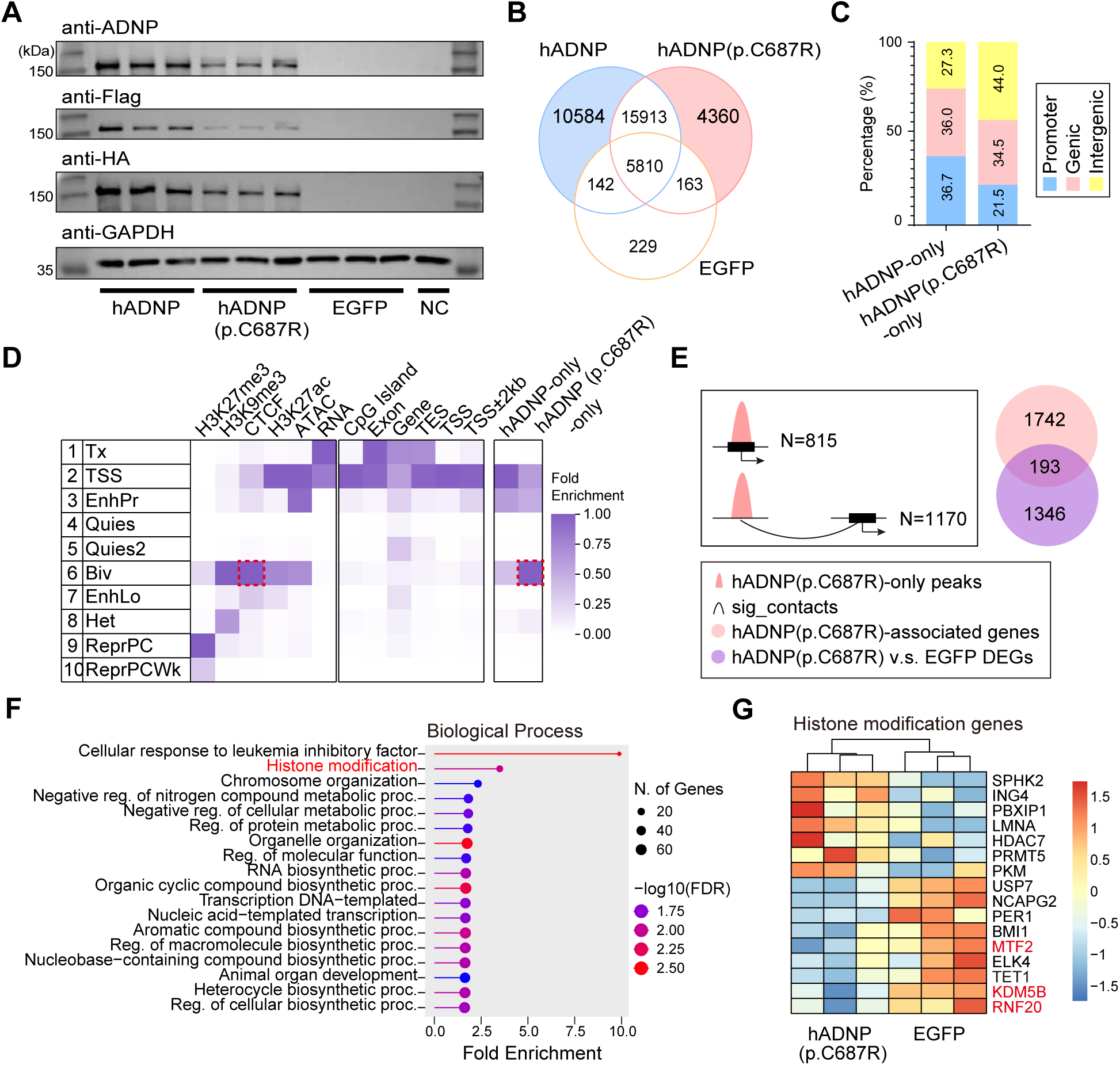
Altered chromatin binding profile of hADNP (p.C687R) in HEK293T cells. (**A**) Western blots showing the expression of hADNP, hADNP (p.C687R), and EGFP in HEK293T cells. NC, untreated cells as negative control. Three antibodies were used: anti- ADNP; anti-HA (N-terminal tag); anti-Flag (C-terminal tag). GAPDH was used as loading control. (**B**) Venn diagram of CUT&Tag peaks among hADNP, hADNP (p.C687R), and EGFP. (**C**) Genomic annotation of unique peaks for ‘hADNP-only’ and ‘hADNP (p.C687R)- only’. Promoter (blue), genic (pink), intergenic (yellow). (**D**) ChromHMM analysis; red dashed squares highlight ‘hADNP (p.C687R)-only’ enrichment in CTCF-associated state #6. (**E**) Left, schematic diagram illustrating identification of genes located directly at ‘hADNP (p.C687R)-only’ peaks and genes connected via distal chromatin interactions. Right, overlap between identified genes and DEGs in hADNP (p.C687R)-overexpressing cells. (**F**) GO analysis of 193 overlapping genes. (**G**) Heatmap of genes in histone modification pathway; methylation-related genes highlighted in red.

Using this system, we performed anti-Flag CUT&Tag to profile genome-wide binding patterns. The clustered correlation heatmap and volcano plots showed clear separation and difference of both hADNP and hADNP (p.C687R) samples from the EGFP control group. Although hADNP and hADNP (p.C687R) clustered closely together, the separation was still evident between the two **(Fig. S5A, S5B)**. Peak overlapping analysis further identified unique binding sites for each protein, with 10,584 sites specific to hADNP (“hADNP_only”) and 4,360 sites specific to hADNP (p.C687R) (“hADNP (p.C687R)_only”) **(Fig. 4B)**. Genomic annotation showed that only 21.5% of hADNP (p.C687R)_only peaks were located at promoters **(Fig. 4C; Table S1)**, suggesting these sites act predominantly through distal regulatory mechanisms.

To characterize chromatin context, we integrated ChromHMM segmentation (Roadmap Epigenomics *et al*, 2015; van der Velde *et al*, 2021) with ChIP-seq and ATAC-seq data from wild-type HEK293T cells. hADNP_only peaks were enriched at TSS state (state #2), with strong active H3K27ac and ATAC-seq signals, whereas hADNP (p.C687R)_only peaks were enriched at bivalent chromatin (state #6), marked by both repressive H3K9me3 and active H3K27ac signal **(Fig. 4D)**. Notably, CTCF, a factor mediating long-range chromatin loops, was also highly enriched at state #6, further suggesting potential distal regulatory activity of hADNP (p.C687R)_only peaks. Using published Hi-C data from wild-type HEK293T cells (Wen *et al*, 2018; Yang *et al*, 2023; Yang *et al*, 2021), we identified genes associated with hADNP (p.C687R)_only peaks either directly or via significant chromatin contacts, defining “hADNP (p.C687R)-associated genes” **(Fig. 4E; Table S2)**.

RNA-seq analysis of the same overexpression system revealed distinct transcriptional profiles among EGFP, hADNP, and hADNP (p.C687R) groups **(Fig. S5C)**. Both hADNP and hADNP (p.C687R) induced substantial differential gene expression compared to EGFP **(Fig. S5D; Table S5-6)**. Among 193 differentially expressed genes overlapping hADNP (p.C687R)-associated genes, GO analysis showed significant enrichment for “histone modification” **(Fig. 4F)**, including key regulators such as MTF2, KDM5B, and RNF20 **(Fig. 4G)**, which are associated with H3K4me3 and H3K27me3 modifications. In contrast, 746 hADNP-associated genes were identified using the same strategy, showing enrichment for cell cycle and metabolic processes **(Fig. S5E-F; Table S3-4)**.

### Altered bivalent chromatin in hADNP (p.C687R) iPSCs independent of transcription

To examine the molecular impact of hADNP (p.C687R) in HVDAS, we generated iPSCs from a patient carrying the mutation and a healthy control. hADNP (p.C687R) (mutant) iPSCs displayed typical pluripotent morphology and expressed OCT4 and NANOG at day 0, but formed abnormally shaped embryoid bodies (EBs) at day 6 during neural induction. Despite these early abnormalities, the mutant line differentiated into NPCs by day 26 **(Fig. 5A)**.

**Figure 5.**
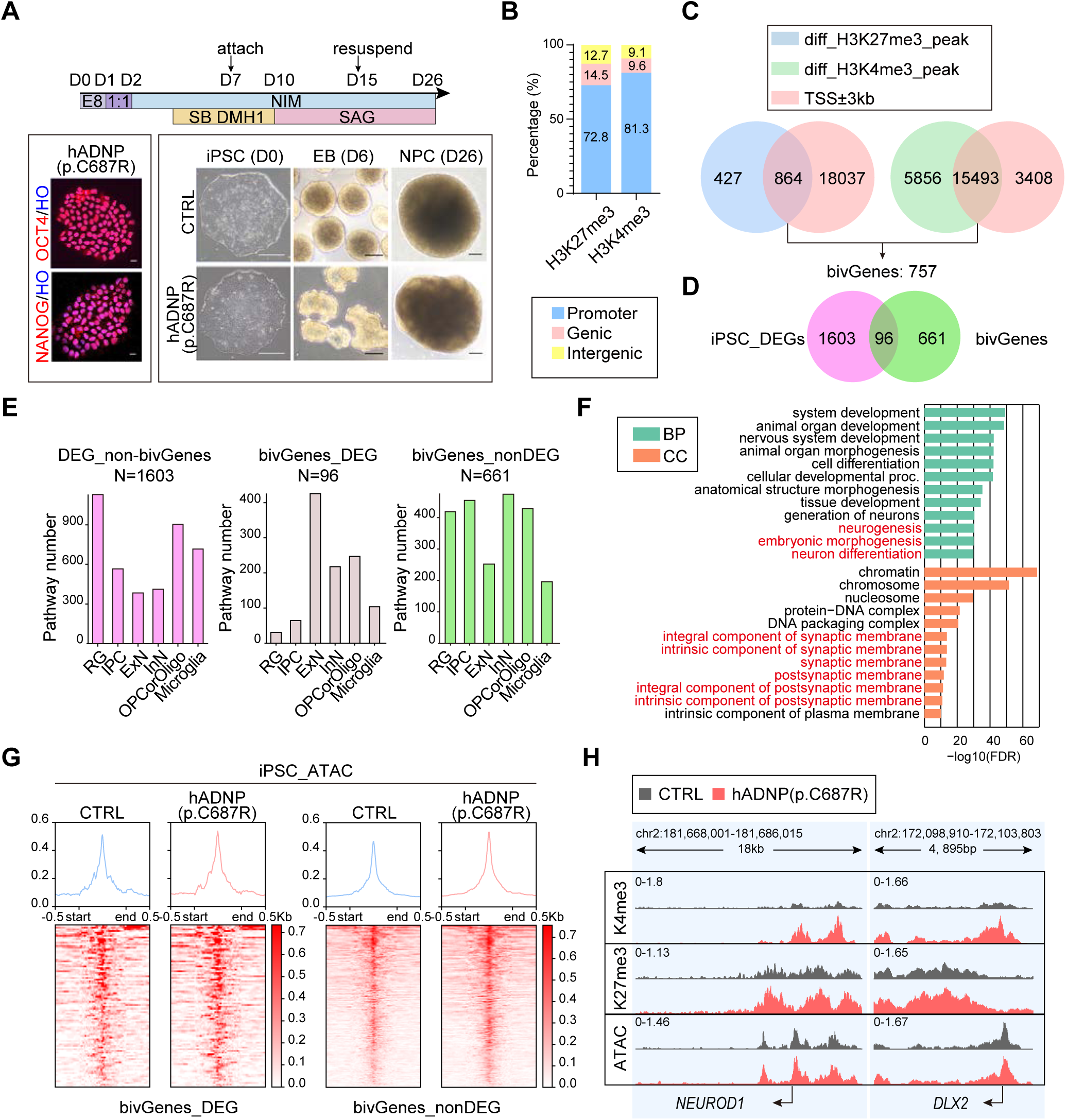
Altered bivalent chromatin in hADNP (p.C687R) iPSCs independent of transcription. (**A**) Schematic of PBMC reprogramming to NPCs. Stem cell markers at Day 0: OCT4/NANOG (red), Hoechst (blue). Scale bars: left IF, 20 μm; right D0, 250 μm; D6, 100 μm; D26, 250 μm. (**B**) Genomic annotation of differential H3K27me3/H3K4me3 ChIP-seq peaks. (**C**) Venn diagrams defining bivalent genes (bivGenes, N = 757) from H3K4me3/H3K27me3 overlaps. (**D**) Overlap between bivGenes and DEGs from iPSC RNA-seq. (**E**) CellGO enrichment of cell type–associated pathways. (**F**) GO analysis of bivGenes_nonDEG (N = 661). (**G**) Aggregated ATAC-seq signals at promoters (TSS ± 3 kb) of bivGenes_DEG and bivGenes_nonDEG. (**H**) IGV tracks of *NEUROD1* and *DLX2*: H3K4me3, H3K27me3, ATAC-seq.

Given the selective effects of hADNP (p.C687R) on genes associated with histone modifications **(Fig. 4F-G)**, we hypothesized that hADNP (p.C687R) disrupts early bivalent chromatin regulation at the stem cell stage. ChIP-seq for the histone marks H3K4me3 and H3K27me3 revealed distinct clustering between mutant and control iPSCs **(Fig. S6A, C)** and global increases in both marks in mutant iPSCs **(Fig. S6B, D; Table S7-8)**. Differential peaks were predominantly located at promoter regions (TSS ±3 Kb) **(Fig. 5B)**. By identifying loci with concurrent changes in H3K4me3 and H3K27me3 at promoter we defined 757 genes with altered bivalent chromatin (bivGenes) **(Fig. 5C)**.

RNA-seq of the same samples revealed transcriptional differences between mutant and control iPSCs **(Fig. S6E–F; Table S9)**, yet most bivGenes were not differentially expressed **(Fig. 5D)**, suggesting that bivalent chromatin alterations occur independently of gene transcription at this stage. CellGO analysis showed that non-bivalent DEGs (DEG_non-bivGenes, N = 1,603) were enriched in radial glial (RG) pathways, while differentially expressed bivGenes (bivGenes_DEG, N = 96) were associated with excitatory neuron (ExN) programs **(Fig. 5E)**. Notably, non-differentially expressed bivGenes (bivGenes_nonDEG, N = 661) were more enriched in inhibitory neuron (InN)– related pathways **(Fig. 5E)** and pathways for neural development and synaptic function **(Fig. 5F)**.

ATAC-seq further revealed distinct chromatin accessibility profiles, with 1,544 upregulated and 281 downregulated peaks in mutant iPSCs **(Fig. S6G–H; Table S10)**, consistent with disrupted chromatin repression of normal hADNP. Integration with the bivalent gene sets showed that both bivGenes_DEG and bivGenes_nonDEG loci resided in open chromatin, with slightly increased accessibility at bivGenes_nonDEG regions in mutant iPSCs **(Fig. 5G)**, which suggest that bivalent chromatin domains in hADNP (p.C687R) cells might be epigenetically primed for activation during later stages of neural differentiation.

Representative IGV tracks of *NEUROD1* and *DLX1*, key regulators of neuronal lineages, illustrate this poised bivalent state **(Fig. 5H)**.

### Activation of BivGenes in hADNP (p.C687R) NPCs associated with GABAergic lineage programs

The identification of poised bivalent genes (BivGenes) in hADNP (p.C687R) iPSCs **(Fig. 5)** prompted us to examine whether these loci become activated during neural differentiation and contribute to altered transcriptional programs. To explore the role of ADNP in neuronal gene regulation, we first generated ADNP-knockout (KO) HEK293T cells using CRISPR/Cas9 **(Fig. S7A–B; Supplement file)**. RNA-seq revealed distinct clustering between WT and KO cells, with 511 genes upregulated and 399 downregulated **(Fig. S7C–D; Table S11)**. Notably, upregulated genes were enriched for synapse-related functions even in this non-neuronal context **(Fig. S7E)** and showed preferential expression in GABAergic neurons **(Fig. S7F–G)**, including increased expression of genes critical for GABAergic development (*GAD1*, *NKX2-1*, *SLC6A1*) **(Fig. S7H)**. These results indicate that ADNP normally represses GABAergic gene programs.

Guided by these findings, we differentiated iPSCs toward a GABAergic fate using SAG and collected NPCs at day 26 **(Fig. 5A)**. ATAC-seq analysis revealed increased chromatin accessibility in hADNP (p.C687R) NPCs **(Fig. S8A–B; Table S12)**, particularly at previously identified BivGene_nonDEG regions **(Fig. 6A)**. Motif analysis of these open regions showed enrichment of NKX family TFs, especially NKX2-1 **(Fig. 6B)**, a key regulator of GABAergic lineage specification. RNA-seq of NPCs demonstrated clear transcriptional divergence between mutant and control cells **(Fig. S8C–D; Table S13)**. Upregulated genes were enriched in neurodevelopmental, synaptic, and axonal pathways, including GABAergic synapses **(Fig. S8E)**. Notably, 26% of BivGenes_nonDEGs (N = 174) became differentially expressed in NPCs **(Fig. 6C)**, forming a BivGene_NPC_DEG subset strongly linked to neurogenesis, neuron fate commitment and InN **(Fig. 6D, Fig S8F)**. Among these, genes critical for GABAergic interneuron differentiation, including *NKX2-1*, *GAD2*, *LHX*, and *DLX* family members, were significantly upregulated in mutant NPCs **(Fig. 6E–F, Fig. S8G)**. Immunostaining further confirmed elevated NKX2-1⁺ and reduced SOX2⁺ cells in mutant NPCs **(Fig. 6G–H)**.

**Figure 6.**
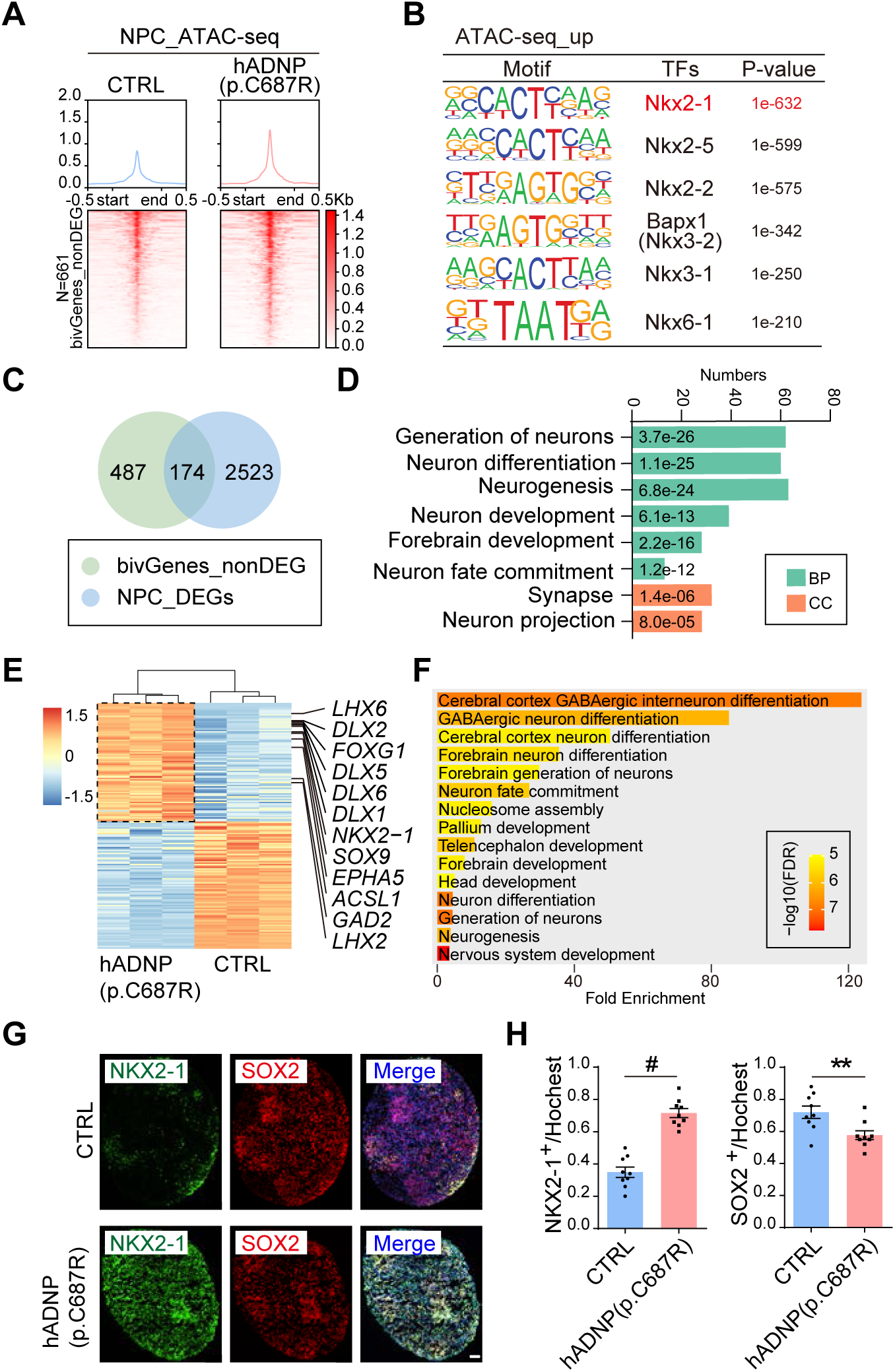
Activation of bivGenes in GABAergic NPCs from hADNP (p.C687R) iPSCs. (**A**) Aggregated ATAC-seq at promoters (TSS ± 3 kb) of bivGenes_nonDEG. (**B**) Predicted TFs at upregulated ATAC-seq peaks (ATAC-seq_up). (**C**) Overlap of bivGenes_nonDEG with DEGs from NPC RNA-seq. (**D**) GO enrichment of 174 overlapping genes (bivGenes_NPC_DEG). (**E**) Heatmap of bivGenes_NPC_DEG, sorted by fold change. (**F**) GO enrichment of upregulated bivGenes_NPC_DEG (N = 84). (**G**) Representative immunofluorescence images: NKX2-1 (green), SOX2 (red), Hoechst (blue). Scale bars, 50 μm. (**H**) Quantification of NKX2-1⁺ and SOX2⁺ cells in (G), normalized to Hoechst⁺ (N = 3/group). Unpaired t-test, ** *P* < 0.01, ^#^ *P* < 0.0001.

Together, these findings indicate that poised bivalent genes become activated in NPCs carrying the ADNP p.C687R mutation and are strongly associated with GABAergic lineage programs, potentially biasing early neural lineage specification in HVDAS.

## Discussion

In summary, our study revisits the pathogenic spectrum of *ADNP* variants and identifies the missense mutation p.C687R as a functionally disruptive allele predicted to alter the ninth ZF domain. *In vitro*, this mutation affects the nuclear localization of ADNP and disrupts the characteristic puncta pattern of the wild-type protein, while *in vivo*, it leads to impaired cortical migration and abnormal neuronal branching. Genomic profiling further revealed mutant-specific chromatin binding and dysregulation of histone modification– related genes. In patient-derived iPSCs, the mutation induced aberrant bivalent chromatin states that prime GABAergic lineage genes for activation, providing mechanistic insight into its pathogenic contribution to HVDAS.

### Functional and mechanistic insights into the ADNP p.C687R mutation

In our current study, we provide multiple lines of evidence indicating that the ADNP missense mutation p.C687R disrupts neuronal development through a gain-of-function (GOF) mechanism. To our knowledge, this represents the first case showing evidence supporting a GOF pathogenic mechanism in ADNP-related neurodevelopmental disorders. In contrast, existing literature has primarily attributed the pathogenesis of HVDAS to dosage insufficiency or a LOF mechanism. Heterozygous microdeletions encompassing ADNP consistently result in HVDAS phenotypes (Huynh *et al*, 2018). Most pathogenic variants identified to date are protein-truncating mutations that lead to the loss of critical C-terminal domains, including NLS, homeobox domain, and HP1-binding regions, all essential for ADNP’s nuclear localization and chromatin regulatory functions (Cappuyns *et al*., 2018; Yan *et al*., 2022). Truncated mutant proteins are typically undetectable in the patient tissues carrying common mutations such as p.C719* (D’Incal *et al*, 2024; Yan *et al*., 2022), suggesting rapid degradation or failed translation. Some missense mutations have also been linked to aberrant splicing, generating truncated products and further reinforcing the LOF model (Georget *et al*, 2023). Additionally, missense mutations within the NAP peptide domain have been proposed to impair ADNP’s neuroprotective function, again consistent with LOF effects (Gozes & Shazman, 2023). Despite the dominance of the LOF hypothesis, the potential for gain-of-toxic-function effects in ADNP mutations has remained largely unexplored. In this study, focusing on the p.C687R variant, we present evidence supporting a distinct GOF mechanism. Structural modeling suggests that p.C687R disrupts the last ZF domain, critical for chromatin binding. Consistent with this, chromatin binding assays show that the mutant protein associates with a distinct set of genomic loci, and transcriptomic analyses reveal dysregulation of genes involved in histone modification. Functional studies further demonstrate the pathogenic impact of this mutation. In HEK293 cells, overexpression of p.C687R alters the subnuclear localization of the mutant protein and disrupts the distribution of co-expressed wild-type ADNP, indicating a dominant-negative effect. *In vivo*, expression of the mutant protein leads to aberrant neuronal migration and disorganized branching during neurodevelopment. However, due to the lack of anti-ADNP antibodies that can distinguish between wild-type and mutant proteins with single amino acid substitutions, it remains technically difficult to directly demonstrate mutant-specific chromatin binding in patient-derived cells. To further validate the GOF mechanism *in vivo*, generating a knock-in mouse model carrying the p.C687R mutation would be a critical next step. A homozygous model, if viable, could provide a definitive system to dissect the functional impact of the mutant protein relative to wild-type ADNP, offering important mechanistic insights.

### Epigenetic mechanisms underlying ADNP p.C687R–mediated neuronal dysregulation

At the molecular level, our findings indicate that the ADNP p.C687R mutation disrupts neuronal differentiation by perturbing chromatin regulation. ADNP is a central chromatin regulator, influencing both transcriptionally active and repressive regions through histone modifications. In mouse models, ADNP haploinsufficiency reduces H3K79me2 (Hadar *et al*, 2021), a mark associated with transcription elongation. A relationship between ADNP and H3K9me3 has been implied. ADNP is recruited to H3K9me3-enriched pericentromeric heterochromatin by interacting with HP1 and represses the satellite repeats (Mosch *et al*, 2011). In addition, a mouse model harboring ADNP-C-terminal-mutation exhibits a decrease in H3K9me2/3, which can be expanded by inhibiting the histone demethylase LSD1, hinting at a regulatory circuit between ADNP and H3K9 methylation (Lin *et al*, 2025). In our overexpression system, p.C687R exhibited a distinct chromatin-binding profile, enriched at loci marked by both H3K9me3 and H3K27ac, and specifically altered the expression of histone-modifying genes. This demonstrates that even a single amino acid change can disrupt ADNP-mediated epigenetic regulation, reshaping chromatin accessibility and transcriptional programs.

Other than affecting histone modifications, ADNP also acts in the form of chromatin remodeling complexes by interacting with other proteins, of which the two most central proteins are HP1 and BRG1. CHAHP complex, composed by ADNP, HP1 and CHD4, participates in the establishment of local inaccessibility in euchromatin and restricts the expression of lineage-specifying genes (Ostapcuk *et al*., 2018). On the other hand, ADNP interacts with some core subunits of the chromatin remodeling SWI/SNF complex, such as BRG1, which utilizes ATP to mediate nucleosome configuration and render DNA accessibility to enhancer or promoter (Mandel & Gozes, 2007). In addition, ADNP, BRG1 and CHD4 also form a complex, which regulates the primitive endoderm (PrE) genes during the early development of mouse embryos. Intriguingly, the repressive effect of ADNP-BRG1-CHD4 complex involves the balance of bivalent-histone markers, H3K27me3 and H3K4me3, which is a special mechanism in developmental genes in embryonic stem cells (ESCs), assuring the rapid priming during differentiation. The H3K4me3/H3K27me3 ratio increased in promoters of PrE genes upon disruption of the ADNP-BRG1-CHD4 complex, which results in the abnormal up-regulation (Sun *et al*, 2020b). Consistent with these mechanisms, in our current study, patient-derived iPSCs carrying p.C687R showed altered bivalent marks at promoters of neurodevelopmental genes. A subset of these genes became transcriptionally activated during differentiation into GABAergic progenitors, indicating that the mutant releases normally repressed developmental programs and biases neuronal lineage specification.

### Role of ADNP in GABAergic neuronal differentiation

While earlier studies have primarily focused on the role of ADNP in excitatory neurons, our findings significantly expand this view by demonstrating its critical involvement in GABAergic neuronal development and function. Conditional deletion of *Adnp* in the dorsal telencephalon during embryogenesis results in a substantial loss of upper-layer cortical neurons (Clemot-Dupont *et al*., 2025), highlighting its essential role in excitatory neuronal differentiation. In models of ADNP haploinsufficiency, impaired excitatory synaptic architecture and transmission have been consistently observed, including altered expression and activity of synaptic markers such as VGLUT1 (Sragovich *et al*, 2019) and CaMKIIα (Cho *et al*., 2023). However, accumulating evidence now indicates that ADNP is also indispensable for the development and functional integrity of inhibitory neuronal circuits. Mouse models harboring C-terminal truncating mutations in *Adnp* exhibit deficits in both GABAergic and glutamatergic neurotransmission (Lin *et al*., 2025). Notably, recent work using human iPSC-derived neurons carrying the *ADNP* p.Tyr719* mutation revealed disrupted GABAergic lineage specification and impaired neuronal maturation (Hu *et al*, 2024). In our study, we provide direct mechanistic evidence supporting ADNP’s role in GABAergic neuronal differentiation. Transcriptomic profiling of ADNP-knockout 293T cells revealed upregulation of genes critical for GABAergic identity, including *NKX2-1* and *GAD1*. Similar patterns were observed in patient-derived iPSCs with the *ADNP* p.C687R mutation, as sites enriched *NKX2-1* binding motifs abnormally gained accessibility upon mutation. Integrated chromatin accessibility and transcriptomic analyses, we identified dysregulated gene networks involved in GABAergic neuron differentiation and synaptogenesis, and immunofluorescent staining further confirmed the transition of cellular state. Our findings in iPSCs strengthen the role of ADNP in specifying GABAergic fate.

In parallel with these findings on neuronal development, previous studies have demonstrated that ADNP exerts neuroprotective effects, particularly through its active peptide NAP, which counteracts glutamate-induced excitotoxicity both *in vitro* (Bassan *et al*., 1999) and *in vivo* (Sokolowska *et al*, 2011). ADNP expression is downregulated in response to NMDA-induced excitotoxicity (Teuchner *et al*, 2011), suggesting a feedback mechanism between glutamatergic signaling and ADNP regulation. Importantly, preclinical studies have shown that low-dose ketamine, an NMDA receptor antagonist, increases *Adnp* expression levels (Brown *et al*, 2015). Building on these findings, a small clinical trial explored ketamine as a potential therapeutic strategy for HVDAS. In this study, EEG recordings revealed increased 40-Hz gamma-band inter-trial coherence (ITC) and decreased 20-Hz beta-band ITC following ketamine administration (Kolevzon *et al*, 2022). These neurophysiological signatures are likely associated with enhanced glutamatergic signaling and GABAergic modulation. Given ADNP’s emerging role in regulating excitatory/inhibitory (E/I) balance, ketamine-induced upregulation of ADNP may offer a promising approach to restoring cortical circuit homeostasis in ADNP-related neurodevelopmental disorders.

### Limitations and perspectives

In this study, we investigate the pathogenicity of the rare missense mutation p.C687R and provide compelling molecular and cellular evidence for a GOF as a pathogenic mechanism in HVDAS. However, several limitations should be acknowledged. First, the current findings are largely based on *in vitro* and iPSC-derived systems; *in vivo* validation is still lacking. Establishing a knock-in mouse model carrying the mADNP (p.C686R) mutation will be crucial for confirming the pathogenic mechanism and for assessing behavioral, cellular, and developmental phenotypes in a physiological context. In addition, the molecular details of how this single amino acid substitution alters ADNP-mediated chromatin regulation remain incompletely understood. Specifically, the precise relationship between ADNP mutant–induced histone modification changes and neuronal lineage specification requires further dissection. Finally, as this study was derived from a single patient case, it remains uncertain whether the observed GOF mechanism represents a generalizable pathogenic route or an individual-specific anomaly.

Despite these limitations, our findings introduce a new pathogenic mechanism for ADNP- related disorders. The identification of a gain-of-function effect provides new insight into ADNP’s chromatin regulatory roles and complements the existing loss-of-function framework. Future work using *in vivo* models and multi-omic approaches will be essential to determine how GOF and LOF mutations converge on shared epigenetic pathways and how these alterations disrupt neuronal differentiation and E/I balance. Elucidating these mechanisms will provide critical insights for developing personalized therapeutic strategies aimed at restoring proper ADNP–chromatin regulation and neural circuit homeostasis in HVDAS.

## Materials & Methods

Our study examined both male and female, and similar findings are reported for both sexes.

### Plasmid construction

Full-length human *ADNP* (NM_015339.5) and the p.C687R variant were generated as previously described (Ge *et al*., 2024), with an N-terminal HA tag and C-terminal Flag/Myc tags followed by an IRES-EGFP cassette. The p.C686R variant was introduced using the KOD-Plus Mutagenesis Kit (TOYOBO). All constructs were verified by Sanger sequencing.

### Cell culture and transfection

HEK293T cells were cultured in DMEM supplemented with 10% FBS at 37°C in 5% CO₂. Plasmid transfection was performed using HighGene reagent (Abclonal, RM09014). Cells were collected 72 h post-transfection for immunostaining, CUT&Tag, or Western blot analysis.

### iPSC reprogramming and neural differentiation

Patient-derived iPSCs were reprogrammed from PBMCs using episomal vectors (Addgene #27077, #27078, #27080) as previously described (Mengnan *et al*, 2024). NPCs were generated via embryoid body-based dual-SMAD inhibition using SB431542 and DMH-1, followed by SAG treatment.

### Immunofluorescence staining (IF)

Cells were fixed with 4% paraformaldehyde (PFA), permeabilized with 0.5% Triton X-100, blocked with 10% goat serum, and incubated with primary antibodies against Flag (1:1000) and Myc (1:1000), followed by Alexa Fluor-conjugated secondary antibodies. Nuclei were counterstained with DAPI or Hoechst. Images were acquired using a Nikon AX confocal microscope, and puncta were quantified in Fiji.

### Western blotting

HEK293T lysates were prepared in SDS buffer, resolved by SDS–PAGE, and transferred to nitrocellulose membranes. Membranes were probed with antibodies against HA, Flag, ADNP, and GAPDH, followed by HRP-conjugated secondary antibodies and ECL detection (Tanon).

### IUE

To assess *in vivo* effects of hADNP (p.C687R), plasmids were introduced into E14.5 CD- 1 mouse embryos via IUE (five 50 ms pulses at 45 V with 950 ms intervals). All mouse work was approved by the Animal Care and Use Committee of Shanghai Medical College, Fudan University. Mice were housed at 21 to 24 °C with 45% to 55% humidity under a 12-hour light/dark cycle. All mice were allowed sterile water and food ad libitum. Both sexes were used. Pups were collected at E18.5 or P14. Brains were fixed in 4% PFA, cryosectioned, and immunostained for Flag and GFP/RFP. Neuronal morphology was analyzed by Sholl analysis in Fiji.

### RNA extraction and RNA-seq

Total RNA from HEK293T cells, iPSCs, and NPCs was extracted using the Direct-zol RNA Microprep Kit (ZYMO). Libraries were prepared from 500 ng of total RNA and sequenced on an Illumina NovaSeq 6000 (paired-end 150 bp).

### CUT&Tag

CUT&Tag assays were performed using the NovoNGS CUT&Tag 3.0 High-Sensitivity Kit (Novoprotein, N259-YH01). Approximately 1×10⁵ cells were immobilized with concanavalin A-coated magnetic beads after 10 min incubation at room temperature. Samples were incubated overnight at 4°C with 1 μg anti-Flag antibody (Smart Lifesciences, SLAB0101), followed by 1 h incubation with secondary antibody at room temperature. After pre-binding with protein A/G–Tn5 transposase, tagmentation was initiated by adding MgCl₂ and incubating at 37°C for 1 h. Reactions were terminated with proteinase K digestion, and DNA fragments were purified, amplified, and sequenced on an Illumina NovaSeq 6000 (paired-end 150 bp).

### ChIP-seq

Native ChIP-seq was performed as described (Sun *et al*, 2024) using approximately 3×10⁶ iPSCs. Cells were lysed in NP40 buffer and chromatin was fragmented with MNase at 28°C for 5 min in douncing buffer (10 mM Tris pH 8, 4 mM MgCl₂, 1 mM CaCl₂). Chromatin was incubated with anti-H3K27me3 (Millipore, 07-449) or anti-H3K4me3 (Abcam, ab8580), followed by capture with protein A/G magnetic beads (Thermo Scientific, 88803). Beads were sequentially washed with low-salt, high-salt, LiCl, and TE buffers, and chromatin was eluted with 0.1 M NaHCO₃/1% SDS. After RNase A and proteinase K digestion, DNA was purified by ethanol precipitation and size-selected using SPRI beads (Beckman, B23318). Libraries were prepared using the VAHTS Universal End Preparation and Adapter Ligation Modules (Vazyme, N203-01 and N204-01), followed by amplification with VAHTS HiFi Universal Mix (Vazyme, N618-02), and sequenced on an Illumina NovaSeq (paired-end 150 bp).

### ATAC-seq

ATAC-seq was performed using the TruePrep DNA Library Prep Kit V2 for Illumina (Vazyme, TD501-02) as previously described (Zhang *et al*, 2024). Nuclei from 1×10⁵ iPSCs were extracted with NP40 lysis buffer and incubated with Tn5 transposase at 37°C for 45 min. DNA was purified using the MinElute Gel Extraction Kit (Qiagen, 28604), PCR amplified, purified with SPRIselect beads (Beckman, B23318), and sequenced on an Illumina NovaSeq (paired-end 150 bp; Novogene).

### Bioinformatic analysis

Sequencing quality was assessed using FastQC, and reads were trimmed with Trim Galore. Alignment was performed to the human genome (hg38) using HISAT2 (RNA-seq) or Bowtie2 (ChIP-seq, ATAC-seq, CUT&Tag). Peak calling was performed using MACS2 (narrow peaks) and SICER (broad peaks). Differential analyses were conducted with DESeq2 and DiffBind. Motif enrichment was analyzed using HOMER, and genomic annotation was performed using ChIPseeker and clusterProfiler. Hi-C data were processed using HiC-Pro and JuicerTools as described (Jiang *et al*, 2017).

### Protein structure modeling

The wild-type ADNP structure was predicted using AlphaFold2 (Jumper *et al*, 2021). The p.C687R mutation was modeled with OPUS-Mut (Xu *et al*, 2023), which reconstructs side chains for both the wild-type and mutant proteins. Perturbations in side-chain conformation were quantified, and residues exhibiting substantial deviations were identified as affected. Mutations impacting residues at functional sites were considered functionally disruptive. For additional details, see **Supplementary Material**.

## Acknowledgements

This work was supported by STI2030-Major Projects (2021ZD0203000) (Y.J.), the National Natural Science Foundation of China (No. 32571123, No. 81971272, No.32170601) (Y.J.), the natural Science Foundation of Anhui Province (No. 2308085MH255) (Q.X.), and the academic leaders development program of Children’s Hospital of Fudan University (EKXDPY202306) (Q.X.)

## Authors’ contributions

Y.J., M.X., and Q.X. conceived the ideas and supervised the research; Q.C., X.L., M.W. and D.S. designed and performed the experiments; all the other authors contributed to experiment performance and data analysis; all authors contributed to scientific discussion and manuscript preparation.

## Ethics approval and consent to participate

Human blood collection and iPSC usage and experiments were authorized by Research Ethics Board at Children’s Hospital of Fudan University (No.2024204). Mouse usage and experiments were authorized by Animal Care and Use Committee of Fudan University (FE21154).

## Data availability

All the raw sequencing data were deposited in the Genome Sequence Archive (GSA) at China National Center for Bioinformation (CNCB) under accession number HRA014131 (https://ngdc.cncb.ac.cn/gsa-human/s/8N5c6H17 for reviewer access).

## Expanded view figure legends

**Figure S1. Predicted pathogenicity and structural modeling of hADNP (p.C687R).** (**A**) Predicted pathogenicity in clinical (N = 137) and non-clinical (N = 876) variants. P/LP, pathogenic/likely pathogenic; B/LB, benign/likely benign; VUS, variants of uncertain significance; Conflicting, conflicting interpretations of pathogenicity. (**B**) Proportion of variants in epi-ADNP-1/2. (**C**) Predicted structural impacts on full-length ADNP; ZF domains in green (WT) or purple (**hADNP (p.C687R)**).

**Figure S2. Nuclear localization of hADNP and hADNP (p.C687R) in HEK293T cells. (A)** Representative low-magnification (40x) images of HEK293T cells transfected with hADNP-Flag, hADNP (p.C687R)-Myc, or co-transfected with both plasmids. Red, hADNP- Flag. Green, hADNP (p.C687R)-Myc. Blue, DAPI. Scale bars, 20μm.

**Figure S3. Nuclear localization of mADNP and mADNP (p.C686R) in HEK293T cells. (A**) Representative low-magnification (40x) images of HEK293T cells transfected with mADNP-Flag, mADNP (p.C686R)-Myc, or co-transfected with both plasmids. Red, mADNP-Flag; green, mADNP (p.C686R)-Myc; blue, DAPI. Scale bar, 20 μm. (**B**) High-magnification with co-localization analysis (white dashed lines indicating measured path). Scale bar, 2 μm. (**C**) Proportion of cells with different puncta patterns: no puncta, large (>10 μm), small (<10 μm) (Mean ± SEM, 1-2 slices/sample, N = 4 smples/group). (**D**) Violin plots show the number of puncta per cell (Mean ± SEM, 1-2 slices/sample, N = 4 smples/group). Two-tailed Mann–Whitney U test, ^#^*P* < 0.0001. (**E**) Distribution of puncta number per cell (Mean ± SEM, 1-2 slices/sample, N = 4 smples/group). (**F**) Puncta width (>10 μm or <10 μm) frequency.

**Figure S4. Full Western blots of hADNP and hADNP (p.C687R) in HEK293T cells.** Cells were transfected with hADNP, hADNP (p.C687R), or EGFP. NC, untreated cells as a negative control. (**A**) anti-ADNP; (**B**) anti-HA (N-terminal tag); (**C**) anti-Flag (C-terminal tag); (**D**) anti-GAPDH.

**Figure S5. The chromatin binding and transcriptional profile of hADNP and hADNP (p.C687R) in HEK293T cells.** (**A–B**) CUT&Tag: correlation heatmap and volcano plot (N = 3/group; FDR < 0.05; |Log2FC| ≥ 0.585). (**C–D**) RNA-seq PCA and MA plot (N = 3/group; *P* < 0.05; |Log2FC| > 0). (**E**) Up, identification of genes directly bound by ‘hADNP-only’ peaks and genes connected via distal chromatin interactions. Bottom, overlap between identified genes and DEGs in hADNP-overexpressing cells. (**F**) GO enrichment of 746 overlapping genes.

**Figure S6. Epigenomic and transcriptomic profiling of hADNP (p.C687R) iPSCs.** (**A– B**) PCA and volcano plots of H3K27me3 ChIP-seq. N = 3/group, *P* < 0.05, |Log2FC| ≥ 0.585. (**C–D**) PCA and volcano plots of H3K4me3 ChIP-seq. N = 2 CTRL/3 hADNP (p.C687R), FDR < 0.05, |Log2FC| ≥ 0.585. (**E–F**) PCA and MA plot of RNA-seq. N = 3/group; *Padj* < 0.05; |Log2FC| ≥ 0.585. (**G–H**) PCA and volcano plot of ATAC-seq. N = 3/group, *P* < 0.05, |Log2FC| ≥ 0.585.

**Figure S7. Loss of ADNP in HEK293T upregulates GABAergic neuronal genes.** (**A**) CRISPR/Cas9 knockout schematic; gRNAs indicated; red star, stop codon. (**B**) Western blot showing ADNP expression in positive knockout clones (KO), negative non-targeting clones (WT), and untreated control cells (293T). C5C9 was used as a loading control. (**C– D**) RNA-seq PCA and MA plot. N = 3/group; *Padj* < 0.05; |Log2FC| ≥ 1. (**E–G**) ShinyGo analysis. (**E**), cellular component; (**F**), co-expression (Azimuth cell type); (**G**), and CellGO pathway enrichment analysis for upregulated genes. (**H**) Representative GABAergic markers (*GAD1*, *NKX2-1*, *SLC6A1*) in WT vs. KO. N = 3/group; ^#^ *Padj* < 0.0001.

**Figure S8. Chromatin accessibility and RNA-seq profiling in GABAergic NPCs derived from hADNP (p.C687R) patient iPSCs.** (**A**) ATAC-seq correlation heatmap in hADNP (p.C687R) vs. CTRL NPCs. (**B**) Differential ATAC-seq volcano plot. N = 3/group; FDR < 0.05; |Log2FC| ≥ 0.585. (**C–D**) RNA-seq PCA and MA plot. N = 3/group; *Padj* < 0.05; |Log2FC| ≥ 0.585. (**E**) GO enrichment of upregulated genes. (**F**) CellGO enrichment of bivGenes_NPC_DEG. (**G**) Expression of representative *NKX* and *DLX* genes. N = 3/group; ^#^ *Padj* < 0.0001.

## References

1. Bassan M, Zamostiano R, Davidson A, Pinhasov A, Giladi E, Perl O, Bassan H, Blat C, Gibney G, Glazner G et al (1999) Complete sequence of a novel protein containing a femtomolar-activity-dependent neuroprotective peptide. J Neurochem 72: 1283–1293

2. Bend EG, Aref-Eshghi E, Everman DB, Rogers RC, Cathey SS, Prijoles EJ, Lyons MJ, Davis H, Clarkson K, Gripp KW et al (2019) Gene domain-specific DNA methylation episignatures highlight distinct molecular entities of ADNP syndrome. Clin Epigenetics 11: 64

3. Brown BP, Kang SC, Gawelek K, Zacharias RA, Anderson SR, Turner CP, Morris JK (2015) In vivo and in vitro ketamine exposure exhibits a dose-dependent induction of activity-dependent neuroprotective protein in rat neurons. Neuroscience 290: 31–40

4. Cappuyns E, Huyghebaert J, Vandeweyer G, Kooy RF (2018) Mutations in ADNP affect expression and subcellular localization of the protein. Cell Cycle 17: 1068–1075

5. Chen S, Charness ME (2008) Ethanol inhibits neuronal differentiation by disrupting activity-dependent neuroprotective protein signaling. Proc Natl Acad Sci U S A 105: 19962–19967

6. Cho H, Yoo T, Moon H, Kang H, Yang Y, Kang M, Yang E, Lee D, Hwang D, Kim H et al (2023) Adnp- mutant mice with cognitive inflexibility, CaMKIIalpha hyperactivity, and synaptic plasticity deficits. Mol Psychiatry 28: 3548–3562

7. Clemot-Dupont S, Lourenco Fernandes JA, Larrigan S, Sun X, Medisetti S, Stanley R, El Hankouri Z, Joshi SV, Picketts DJ, Shekhar K et al (2025) The chromatin remodeler ADNP regulates neurodevelopmental disorder risk genes and neocortical neurogenesis. Proc Natl Acad Sci U S A 122: e2405981122

8. D’Incal CP, Cappuyns E, Choukri K, De Man K, Szrama K, Konings A, Bastini L, Van Meel K, Buys A, Gabriele M et al (2024) Tracing the invisible mutant ADNP protein in Helsmoortel-Van der Aa syndrome patients. Sci Rep 14: 14710

9. Ganaiem M, Karmon G, Ivashko-Pachima Y, Gozes I (2022) Distinct Impairments Characterizing Different ADNP Mutants Reveal Aberrant Cytoplasmic-Nuclear Crosstalk. Cells 11

10. Ge C, Tian Y, Hu C, Mei L, Li D, Dong P, Zhang Y, Li H, Sun D, Peng W et al (2024) Clinical impact and in vitro characterization of ADNP variants in pediatric patients. Mol Autism 15: 5

11. Georget M, Lejeune E, Buratti J, Servant E, le Guern E, Heron D, Keren B, de Sainte Agathe JM (2023) Loss of function of ADNP by an intragenic inversion. *Eur J Hum Genet* 31: 967-970

12. Gozes I, Shazman S (2023) A novel davunetide (NAPVSIPQQ to NAPVSIPQE) point mutation in activity- dependent neuroprotective protein (ADNP) causes a mild developmental syndrome. Eur J Neurosci 58: 2641–2652

13. Hadar A, Kapitansky O, Ganaiem M, Sragovich S, Lobyntseva A, Giladi E, Yeheskel A, Avitan A, Vatine GD, Gurwitz D et al (2021) Introducing ADNP and SIRT1 as new partners regulating microtubules and histone methylation. Mol Psychiatry 26: 6550–6561

14. Helsmoortel C, Vulto-van Silfhout AT, Coe BP, Vandeweyer G, Rooms L, van den Ende J, Schuurs- Hoeijmakers JH, Marcelis CL, Willemsen MH, Vissers LE et al (2014) A SWI/SNF-related autism syndrome caused by de novo mutations in ADNP. Nat Genet 46: 380–384

15. Hu R, Boshans LL, Zhu B, Cai P, Tao Y, Youssef M, Girrbach GI, Song Y, Wang X, Tsankov A et al (2024) Expanding GABAergic Neuronal Diversity in iPSC-Derived Disease Models. bioRxiv: 2024.2012.2003.626438

16. Huynh MT, Boudry-Labis E, Massard A, Thuillier C, Delobel B, Duban-Bedu B, Vincent-Delorme C (2018) A heterozygous microdeletion of 20q13.13 encompassing ADNP gene in a child with Helsmoortel-van der Aa syndrome. Eur J Hum Genet 26: 1497–1501

17. Ivashko-Pachima Y, Sayas CL, Malishkevich A, Gozes I (2017) ADNP/NAP dramatically increase microtubule end-binding protein-Tau interaction: a novel avenue for protection against tauopathy. Mol Psychiatry 22: 1335–1344

18. Jiang Y, Loh YE, Rajarajan P, Hirayama T, Liao W, Kassim BS, Javidfar B, Hartley BJ, Kleofas L, Park RB et al (2017) The methyltransferase SETDB1 regulates a large neuron-specific topological chromatin domain. Nat Genet 49: 1239–1250

19. Jumper J, Evans R, Pritzel A, Green T, Figurnov M, Ronneberger O, Tunyasuvunakool K, Bates R, Zidek A, Potapenko A et al (2021) Highly accurate protein structure prediction with AlphaFold. Nature 596: 583–589

20. Kaaij LJT, Mohn F, van der Weide RH, de Wit E, Bühler M (2019) The ChAHP Complex Counteracts Chromatin Looping at CTCF Sites that Emerged from SINE Expansions in Mouse. Cell 178: 1437

21. Kolevzon A, Levy T, Barkley S, Bedrosian-Sermone S, Davis M, Foss-Feig J, Halpern D, Keller K, Kostic A, Layton C et al (2022) An open-label study evaluating the safety, behavioral, and electrophysiological outcomes of low-dose ketamine in children with ADNP syndrome. HGG Adv 3: 100138

22. Lin CH, Ren Y, Tam KW, Conrow-Graham M, Yan Z (2025) Synaptic Deficits in Adnp-Mutant Mice Are Ameliorated by Histone Demethylase LSD1 Inhibition. Autism Res 18: 1342–1355

23. Mandel S, Gozes I (2007) Activity-dependent neuroprotective protein constitutes a novel element in the SWI/SNF chromatin remodeling complex. J Biol Chem 282: 34448–34456

24. Mengnan W, Yan C, Qiong X, Man X (2024) Generation of a human induced pluripotent stem cell line (FDIBSi001-A) from a patient with ADNP syndrome carrying ADNP mutation (c. 2059 T>C). Stem Cell Res 81: 103550

25. Merenlender-Wagner A, Malishkevich A, Shemer Z, Udawela M, Gibbons A, Scarr E, Dean B, Levine J, Agam G, Gozes I (2015) Autophagy has a key role in the pathophysiology of schizophrenia. Mol Psychiatr 20: 126–132

26. Mosch K, Franz H, Soeroes S, Singh PB, Fischle W (2011) HP1 recruits activity-dependent neuroprotective protein to H3K9me3 marked pericentromeric heterochromatin for silencing of major satellite repeats. PLoS One 6: e15894

27. Ostapcuk V, Mohn F, Carl SH, Basters A, Hess D, Iesmantavicius V, Lampersberger L, Flemr M, Pandey A, Thoma NH et al (2018) Activity-dependent neuroprotective protein recruits HP1 and CHD4 to control lineage-specifying genes. Nature 557: 739–743

28. Pascolini G, Di Zenzo G, Panebianco A, Didona B, Gozes I (2024) Extended phenotypic characterization of a novel Helsmoortel-van der Aa syndrome case series. Am J Med Genet A 194: e63539

29. Pinhasov A, Mandel S, Torchinsky A, Giladi E, Pittel Z, Goldsweig AM, Servoss SJ, Brenneman DE, Gozes I (2003) Activity-dependent neuroprotective protein: a novel gene essential for brain formation. Brain Res Dev Brain Res 144: 83–90

30. Roadmap Epigenomics C, Kundaje A, Meuleman W, Ernst J, Bilenky M, Yen A, Heravi-Moussavi A, Kheradpour P, Zhang Z, Wang J et al (2015) Integrative analysis of 111 reference human epigenomes. Nature 518: 317–330

31. Shapira G, Karmon G, Hacohen-Kleiman G, Ganaiem M, Shazman S, Theotokis P, Grigoriadis N, Shomron N, Gozes I (2025) ADNP is essential for sex-dependent hippocampal neurogenesis, through male unfolded protein response and female mitochondrial gene regulation. Mol Psychiatr 30: 2696–2706

32. Siper PM, Layton C, Levy T, Lurie S, Benrey N, Zweifach J, Rowe M, Tang L, Guillory S, Halpern D et al (2021) Sensory Reactivity Symptoms Are a Core Feature of ADNP Syndrome Irrespective of Autism Diagnosis. Genes (Basel) 12

33. Sokolowska P, Passemard S, Mok A, Schwendimann L, Gozes I, Gressens P (2011) Neuroprotective effects of NAP against excitotoxic brain damage in the newborn mice: implications for cerebral palsy. Neuroscience 173: 156–168

34. Sragovich S, Malishkevich A, Piontkewitz Y, Giladi E, Touloumi O, Lagoudaki R, Grigoriadis N, Gozes I (2019) The autism/neuroprotection-linked ADNP/NAP regulate the excitatory glutamatergic synapse. Transl Psychiatry 9: 2

35. Sun D, Zhu Y, Peng W, Zheng S, Weng J, Dong S, Li J, Chen Q, Ge C, Liao L et al (2024) SETDB1 regulates short interspersed nuclear elements and chromatin loop organization in mouse neural precursor cells. Genome Biol 25: 175

36. Sun X, Peng X, Cao Y, Zhou Y, Sun Y (2020a) ADNP promotes neural differentiation by modulating Wnt/beta-catenin signaling. Nat Commun 11: 2984

37. Sun X, Yu W, Li L, Sun Y (2020b) ADNP Controls Gene Expression Through Local Chromatin Architecture by Association With BRG1 and CHD4. Front Cell Dev Biol 8: 553

38. Teuchner B, Dimmer A, Humpel C, Amberger A, Fischer-Colbrie R, Nemeth J, Waschek JA, Kieselbach G, Kralinger M, Schmid E et al (2011) VIP, PACAP-38, BDNF and ADNP in NMDA-induced excitotoxicity in the rat retina. Acta Ophthalmol 89: 670-675

39. van der Velde A, Fan K, Tsuji J, Moore JE, Purcaro MJ, Pratt HE, Weng Z (2021) Annotation of chromatin states in 66 complete mouse epigenomes during development. Commun Biol 4: 239

40. Van Dijck A, Vulto-van Silfhout AT, Cappuyns E, van der Werf IM, Mancini GM, Tzschach A, Bernier R, Gozes I, Eichler EE, Romano C et al (2019) Clinical Presentation of a Complex Neurodevelopmental Disorder Caused by Mutations in ADNP. Biol Psychiatry 85: 287–297

41. Wen Z, Huang ZT, Zhang R, Peng C (2018) ZNF143 is a regulator of chromatin loop. Cell Biol Toxicol 34: 471–478

42. Xu G, Wang Q, Ma J (2023) OPUS-Mut: Studying the Effect of Protein Mutation through Side-Chain Modeling. J Chem Theory Comput 19: 1629–1640

43. Yan Q, Wulfridge P, Doherty J, Fernandez-Luna JL, Real PJ, Tang HY, Sarma K (2022) Proximity labeling identifies a repertoire of site-specific R-loop modulators. Nat Commun 13: 53

44. Yang Y, Li C, Chen Z, Zhang Y, Tian Q, Sun M, Zhang S, Yu M, Wang G (2023) An intellectual disability- related MED23 mutation dysregulates gene expression by altering chromatin conformation and enhancer activities. Nucleic Acids Res 51: 2137–2150

45. Yang Y, Ye X, Dai R, Li Z, Zhang Y, Xue W, Zhu Y, Feng D, Qin L, Wang X et al (2021) Phase separation of Epstein-Barr virus EBNA2 protein reorganizes chromatin topology for epigenetic regulation. Commun Biol 4: 967

46. Zemlyak I, Furman S, Brenneman DE, Gozes I (2000) A novel peptide prevents death in enriched neuronal cultures. Regul Pept 96: 39–43

47. Zhang Y, Li J, Zhao Y, Huang Y, Shi Z, Wang H, Cao H, Wang C, Wang Y, Chen D et al (2024) Arresting the bad seed: HDAC3 regulates proliferation of different microglia after ischemic stroke. Sci Adv 10: eade6900

